# Life history of the Rio Grande leopard frog (*Lithobates berlandieri*) in Texas

**DOI:** 10.1101/364141

**Authors:** Daniel F. Hughes, Walter E. Meshaka

## Abstract

We ascertained various life-history traits from an examination of 310 museum specimens of the Rio Grande leopard frog (*Lithobates berlandieri* Baird, 1859) collected during 1907–2016 from Texas, USA. *Lithobates berlandieri* was captured during every month of the year except November, and adults were most frequently encountered during January–September with a distinct peak in May. Mean body size of adult males (69.5 mm) was smaller than that of adult females (77.5 mm), and both sexes were larger in mean body sizes than those of New Mexico populations (M = 64.4 mm; F = 73.5 mm). Females were gravid during January–September, and most gravid females were captured from late-winter to early-summer. Gonadal enlargement in males was generally high throughout January–September with no detectable seasonal increase. Feeding became widespread in both sexes during May–June shortly after a spring breeding bout. Spent females were common in July and lipid deposition increased in June/July, signaling oogenesis for breeding in the fall. From 15 gravid females, we estimated a mean clutch size of 3,107 eggs which was correlated with female body size, yet egg diameter was not related to clutch or body size. Age to metamorphosis was likely 2 to 4 months depending upon whether eggs were laid in the winter/spring or late fall. If metamorphosis occurred in May/June, the minimum size at sexual maturity in adult males (50.1 mm) could have been reached in 3–4 months and in 6–7 months for adult females (57.2 mm). Mean adult body sizes, however, may have taken 12 to 17 months to reach. A synthesis across Texas populations suggests that the breeding season extends almost continuously from the fall through the winter and spring until mid-summer and is interrupted by winter and summer peaks in seasonal temperatures.

## Introduction

The Rio Grande leopard frog (Anura: Ranidae: *Lithobates berlandieri* Baird, 1859) ranges from the extreme southern border of New Mexico to central Texas, USA, and south, mostly along the gulf coast, to the Mexican states of Hidalgo and Veracruz (Platts 1991; Degenhardt et al. 1996; Stebbins 2003; Santos-Barrera et al. 2010; Tipton et al. 2012; Dixon 2013; Powell et al. 2016; Frost 2018). Introduced populations of the Rio Grande leopard frog occur in several areas of North America, including expanding populations in Arizona, California, and Utah (USA: Clarkson and Rorabaugh 1989; Platz et al. 1990; Rorabaugh et al. 2002; Brennan and Holycross 2006; Stebbins and McGinnis 2012) and well-established populations in river drainages from Baja California and Sonora (México: Rorabaugh 2008; Kraus 2009). The distribution of *L*. *berlandieri* in Texas spans from El Paso to Dallas and south to Brownsville (type locality [see Frost 2018]), where it can be found in a range of habitats from deserts to woodlands in association with generally clear waterways, including rivers, springs, and canals, but also temporary tanks (Axtell 1959; Jung et al. 2002; Santos-Barrera et al. 2010; Tipton et al. 2012; Dixon 2013; Davis and LaDuc 2018). Depending upon habitat availability, presence of sympatric species and vertebrate predators, the Rio Grande leopard frog breeds in both streams and ponds (Rorabaugh 2005; Dodd 2013).

The broad geographic range of *L*. *berlandieri* and its complex evolutionary history in the *L*. *pipiens* group that readily hybridizes with related species (Mecham 1969; Hillis 1988), has prompted its inclusion in numerous studies into its phylogenetic position and extent of intergradation with conspecifics (e.g., Hillis 1981, 1982; Frost and Platz 1983; Hillis et al. 1983; Zaldívar-Riverón et al. 2004). Moreover, introductions of *L*. *berlandieri* outside its native range have been well-documented and those populations remain closely followed (Platz 1990; Clarkson and Rorabaugh 1989; Rorabaugh et al. 2002; Rorabaugh 2008; Kraus 2009). Studies focused on natural-history aspects of *L*. *berlandieri*, however, are rare throughout the species’ natural range, with few exceptions (e.g., Jung et al. 2002; Parker and Goldstein 2004). Given the numerous studies of *L*. *berlandieri*, it is curious that we know essentially nothing about its basic life history, demography, or population biology beyond assumed parallels to other leopard frog species (Platz 1991; Degenhardt et al. 1996; Tipton et al. 2012). For example, we currently lack a reported clutch or egg size for *L*. *berlandieri* (Rorabaugh 2005; Dodd 2013). Consequently, geographic variation in life history for *L*. *berlandieri* is a heretofore unstudied subject. The absence of these data are particularly alarming given that climate change is imperiling amphibians worldwide (Jetz and Pyron 2018), and testing life-history predictions to understand how populations will cope with spatially heterogeneous warming requires information on traits such as clutch and egg size (Sheridan et al. 2018). Further, a rigorous examination into the life-history evolution of North American leopard frogs requires reproductive data for each described lineage and, ideally, from multiple populations within a lineage (Miaud et al. 1999).

Geographic variation in life history has important implications for the study of demography, population ecology, and conservation biology. Without data from populations across a species geographic distribution, however, our understanding of the role geography plays in a species’ life history is significantly diminished. *Lithobates berlandieri,* which ranges from the American Southwest to southeastern México, is bereft of life-history information and the limited data that exist are based on a small handful of reports from isolated populations across its extensive geographic distribution. Here, we used museum specimens to investigate several life-history traits of the Rio Grande leopard frog, including age and body size at sexual maturity, breeding season, clutch size, egg size, sexual-size dimorphism, and larval metamorphosis season. We undertook this project with the aim to provide new, and for some traits complementary, descriptive data on the reproductive biology of *L*. *berlandieri* in Texas, USA.

## Materials and Methods

We examined 310 specimens representing nearly all major life-history stages of the Rio Grande leopard frog housed in three US-based collections: Smithsonian National Museum of Natural History (USNM), Washington, D.C. (196 specimens), Carnegie Museum of Natural History (CM), Pittsburgh, Pennsylvania (77 specimens), and UTEP Biodiversity Collections (UTEP), El Paso, Texas (37 specimens). The specimens were collected during a 109-year period (1907–2016) from 39 counties across most of the species’ core range in Texas (Figure 1).

**Figure 1.**
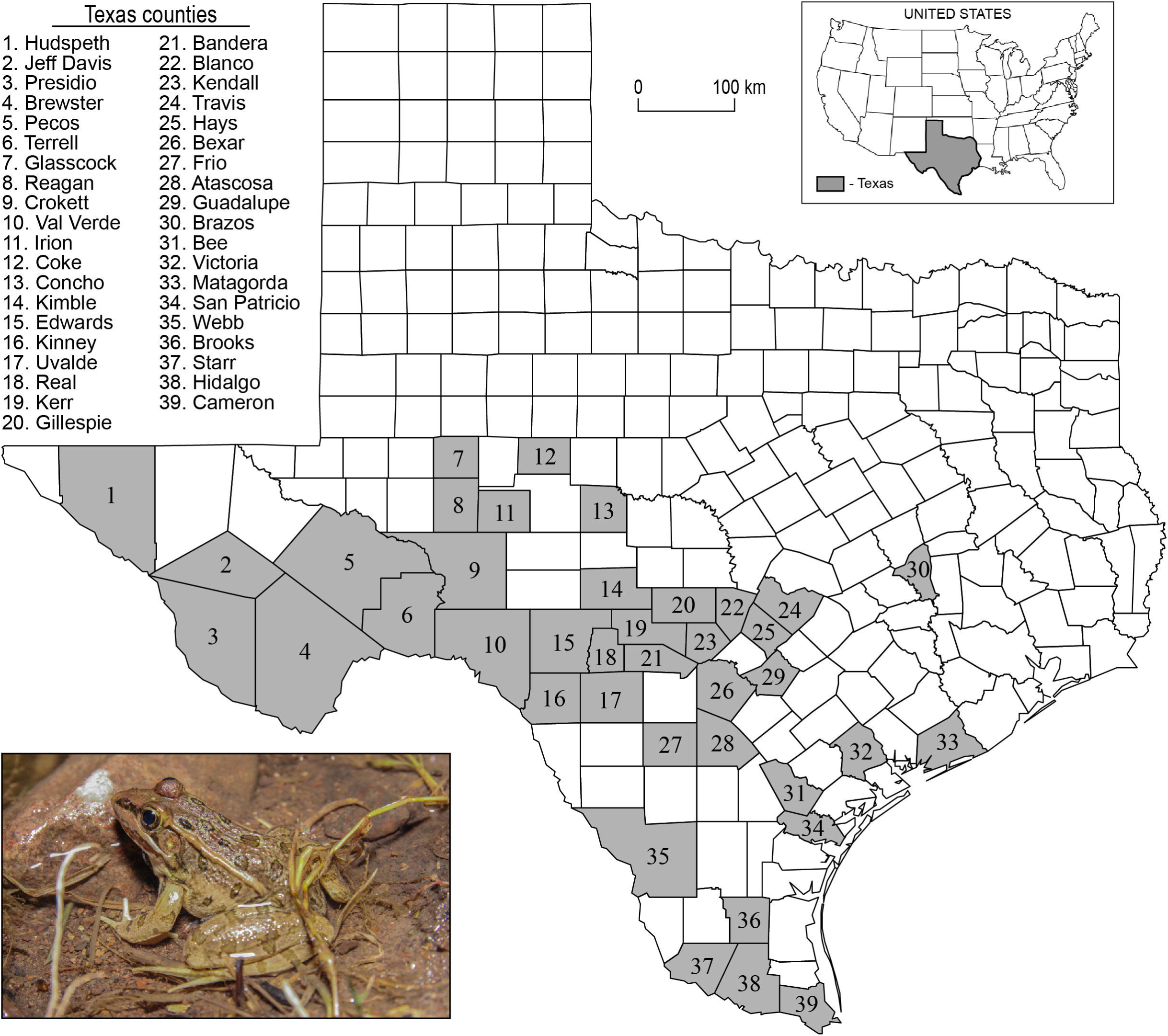
Map of Texas (USA) showing the 39 counties of origin (in gray) for 310 museum specimens of the Rio Grande leopard frog (*Lithobates berlandieri*) examined in this study. Bottom left: Picture of *L*. *berlandieri* from Jeff Davis County (photo by Frank Portillo).

We measured the body size in snout-vent length (SVL) of all specimens to the nearest 0.1 mm using hand calipers. We distinguished recently metamorphosed frogs from tadpoles by the presence of forelimbs (Gosner stage 42) and from juveniles from the presence of a tail stub (Gosner stage 45) (Gosner 1960). We used the presence of swollen testes in combination with enlarged thumbs to determine sexual maturity in males. We measured testis length and mid-width from mature male specimens and estimated male fertility by calculating testis dimensions as a percentage of male SVL. To reduce some of the potential errors derived from preservation artifacts, all reproductive organs of male specimens were measured from the left side of the body (Lee 1982). The incidence of males with enlarged thumbs also served as an estimate of seasonal fertility patterns. We calculated sexual-size dimorphism (SSD) by dividing the mean adult SVL of males by that of females.

We assessed the extent of lipid deposits associated with gonads (i.e., fat bodies) in the body cavity of specimens based on three criteria: 1) trace amounts or no fat bodies; 2) intermediate volume of fat bodies; and 3) high volume of fat bodies that extend anteriorly within the body cavity (Meshaka 2001). We used the highest score as an estimate of the monthly incidence of extensive fat bodies relative to all males and females examined in each month. We recorded the presence of food in the stomach as a proxy for the monthly frequency of individuals that had been feeding for all males or females examined from each month.

As per Meshaka (2001), we assigned sexual maturity in females based on the following four ovarian stages: 1) oviducts were thin and uncoiled, and the ovaries were somewhat opaque; 2) oviducts were larger and more coiled, and the ovaries contained some pigmented oocytes; 3) oviducts were thick and heavily coiled, and the ovaries were in various stages of clutch development; and 4) oviducts were thick and heavily coiled, and the ovaries were full of polarized ova with few non-polarized ova, evidence of a mature clutch and gravid female.

We randomly selected 15 gravid females (ovarian stage 4) to examine for various clutch characteristics. We dissected the clutch out of the body cavity, gently removed excess moisture with a paper towel, and massed the entire clutch to the nearest 0.1 g with an electronic scale. We counted and then massed a subset of mature ova from each clutch ranging 35–100 eggs. We extrapolated the mass of the counted subset of eggs to the weight of the entire brood to provide an estimate of clutch size. We measured the diameters of 10 randomly chosen ova from each of the 15 clutches using an ocular micrometer to the nearest 0.1 mm. We used the largest ovum from each clutch to assess comparative relationships with clutch and female body sizes.

The Texas county map was generated initially using ArcGIS Online and the final figure was prepared with Adobe Illustrator (Creative Cloud: Adobe Systems Inc., San Jose, California, USA). We used Excel 2016 (Microsoft Inc., Redmond, Washington, USA) and the open-access program R version 3.4.4 (R Core Team 2018) to organize numerical data, generate quantitative graphics, and conduct statistical analyses. We present mean measurements followed by ±1 standard deviation (SD). Life-history data was not transformed if it met parametric assumptions, yet in cases when data did not meet assumptions, we used non-parametric tests (Zar 2010). We compared means between samples using two sample t-tests, compared variances using ANOVA F-tests and Levene’s test, and examined relationships between selected variables with correlation coefficients. We recognized statistical significance at *P* < 0.05. A small handful of specimens were either too destroyed for full examination or had had their gastrointestinal tracts, gonads, or both organ systems removed prior to our study, and thus we excluded these specimens from summary statistics.

## Results

### Seasonal incidence of captures

From collection records that span 125 years (1891–2016) for 540 specimens of *L*. *berlandieri* from Texas, we found that individuals were collected in every month except November with a distinct peak in May (n = 165), and only three individuals were captured during October–December (Figure 2A). The sample of 310 specimens we examined mirrored these overall seasonal patterns (Figure 2B). In our sample, specimens were collected during every month of the year except November and December, and only one individual from October (UTEP 12351, an adult male from Hudspeth County [SVL = 63.1 mm]). The interval with the greatest incidence of captured individuals in our sample occurred during January–September, with an apparent peak in captures during late-spring to summer (April–August). Captures suggest corresponding peaks in activity for different life-history classes: Males were captured most during April and May, females in May, and juveniles in July (Figure 2B). Although we examined less than half of the *L*. *berlandieri* specimens at USNM from Texas (196/447 = 44%), the remaining collection records corroborate the paucity of individuals taken during October–December: One record for October in 1899, none in November, and one for December in 1891 (Figure 2A–B).

**Figure 2.**
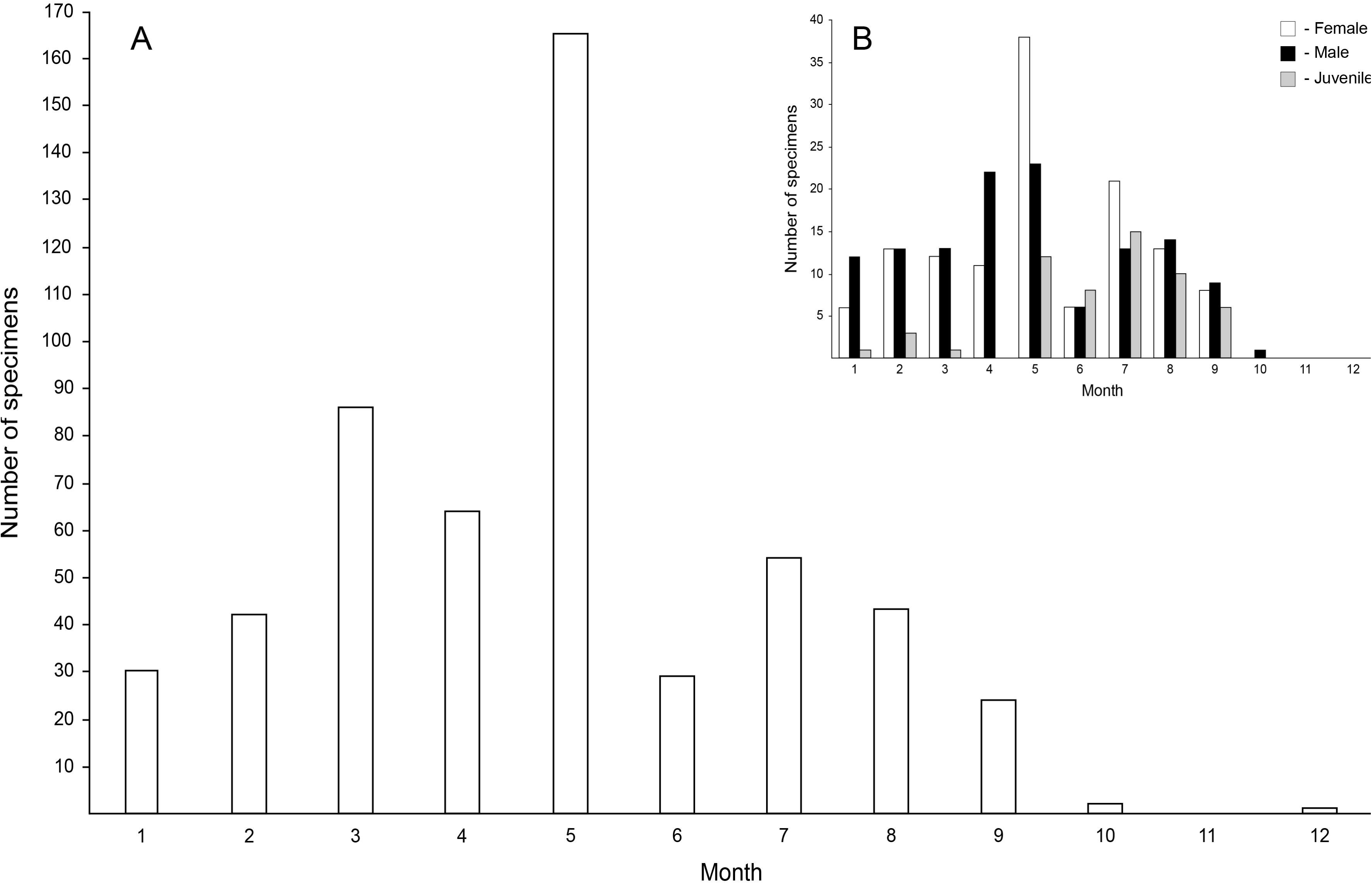
Monthly incidence of captures for 540 specimens (A) and our sample of 310 specimens (B) of the Rio Grande leopard frog (*Lithobates berlandieri*) from Texas (USA). Specimen records originate from three collections: USNM, UTEP, and CM. See text for details on museums.

### Body sizes

The variance in body sizes (SVL) between males (variance = 125.33) and females (variance = 196.25) was not equal (Levene’s test: df = 252, F = 11.5, *P* = 0.00081; F-test: F = 1.57, *P* = 0.0062). The mean SVL of males (69.51 ± 11.19 mm; range 43.6–93.7 mm; n = 126) was significantly smaller than that of females (77.52 ± 14.01 mm; range 47.3–103.2 mm; n = 128) (two sample t-test with unequal variances: t = 5.0333, df = 242, *P* = 0.00000094) (Figure 3). The SSD, expressed as the ratio of mean body sizes of males: females, was 0.897: 1.0. The mean SVL of 43 juveniles was 37.78 ± 10.1 mm (range 18.6–53.8 mm). The mean SVL for 11 specimens that had four legs and a tail was 26.59 ± 3.64 mm (range 19.5–30.9 mm).

**Figure 3.**
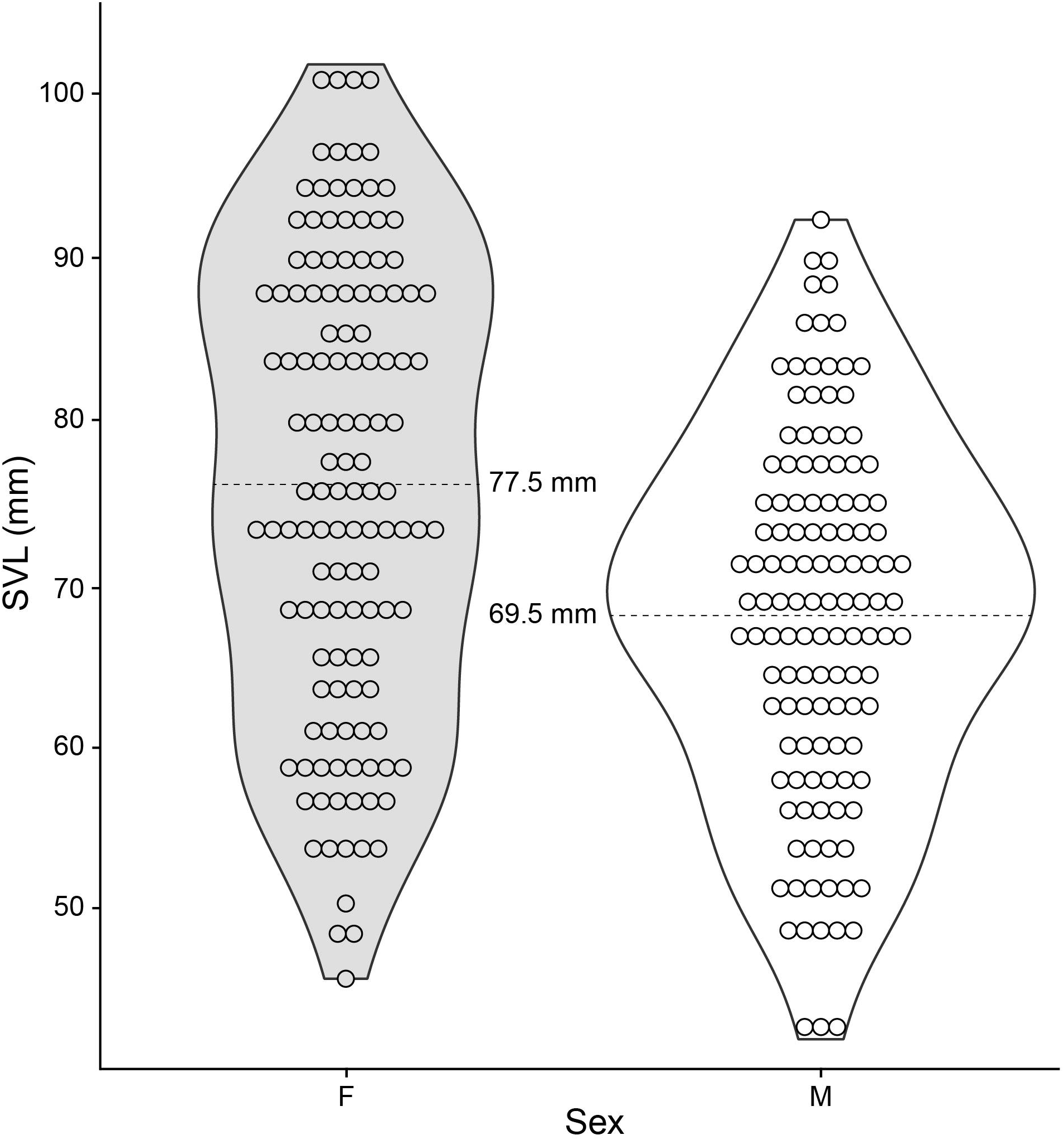
Distribution of body sizes presented as violin plots for 128 female and 126 male Rio Grande leopard frogs (*Lithobates berlandieri*) from Texas (USA). Dashed line indicates the sample mean.

### Male reproduction

Seasonal testis dimensions as a percentage of SVL were generally large during January–September. A seasonal change in testis dimensions was not discernible in our sample (ANOVA: length, F_8,83_ = 0.588, *P* = 0.785; width, F_8,83_ = 1.812, *P* = 0.0863), yet testis length exhibited an apparent increase in late spring (March–May) (Figure 4). Mean testis length as a percentage of male SVL was 8.07 ± 1.5% (range 4.88–12.59%, n = 92) and testis width was 4.11 ± 0.75% (range 2.15–6.40%, n = 92). Nearly 90% of the males in our sample had enlarged thumbs (112/125 males), yet no seasonality to thumb enlargement was detectable.

**Figure 4.**
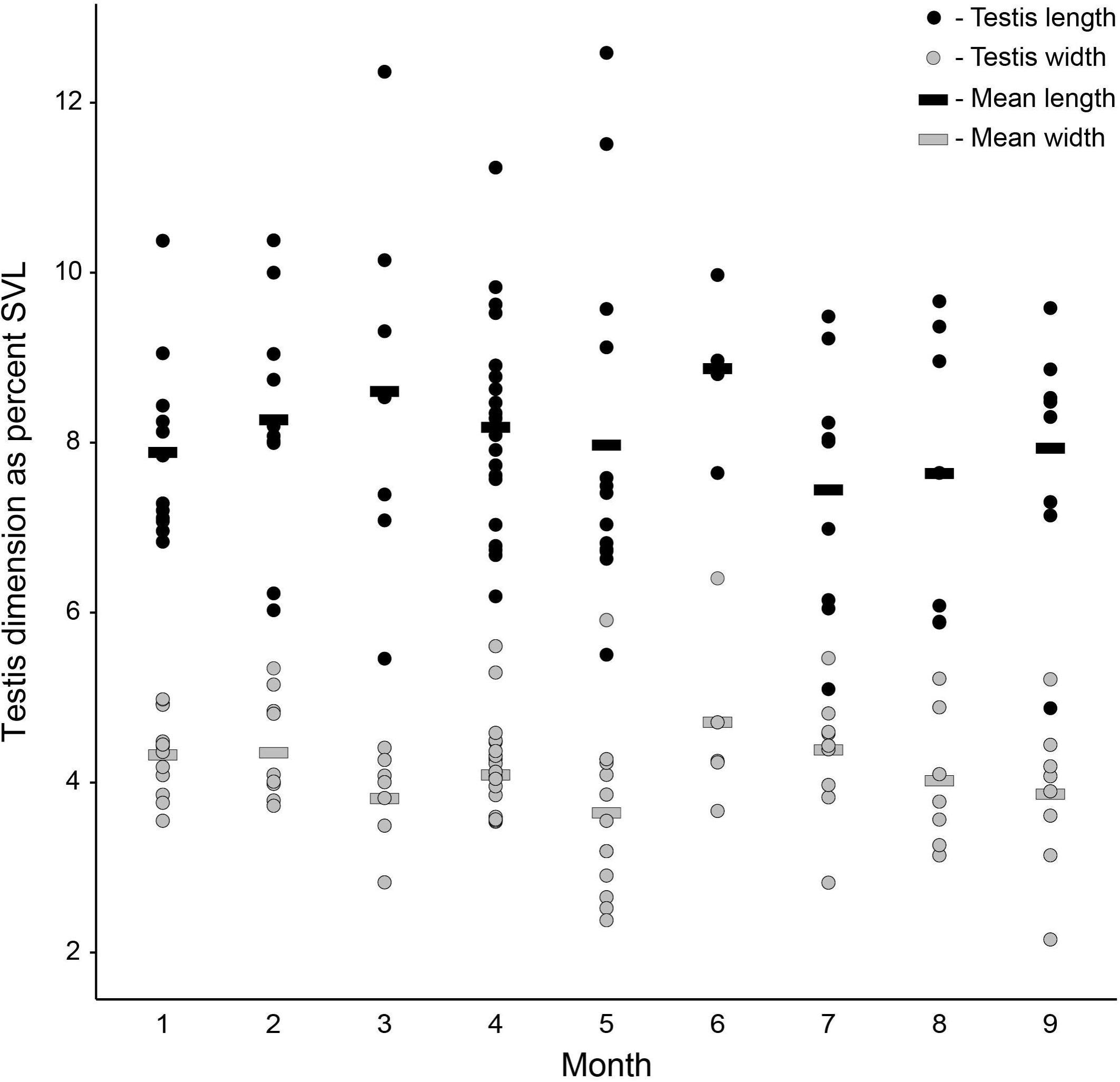
Monthly distribution of testis length and width as a percentage of body size in 92 male Rio Grande leopard frogs (*Lithobates berlandieri*) from Texas (USA).

### Fat and food periodicity

From January–September, the monthly percentage of males with extensive fat development was lowest March–May, and rapidly increased to a peak in July and tapered off slowly (Figure 5A). The monthly percentage of males containing food was lowest in March (0%) and increased rapidly to 100% of the males in May and June, then precipitously dropped (Figure 5A).

**Figure 5.**
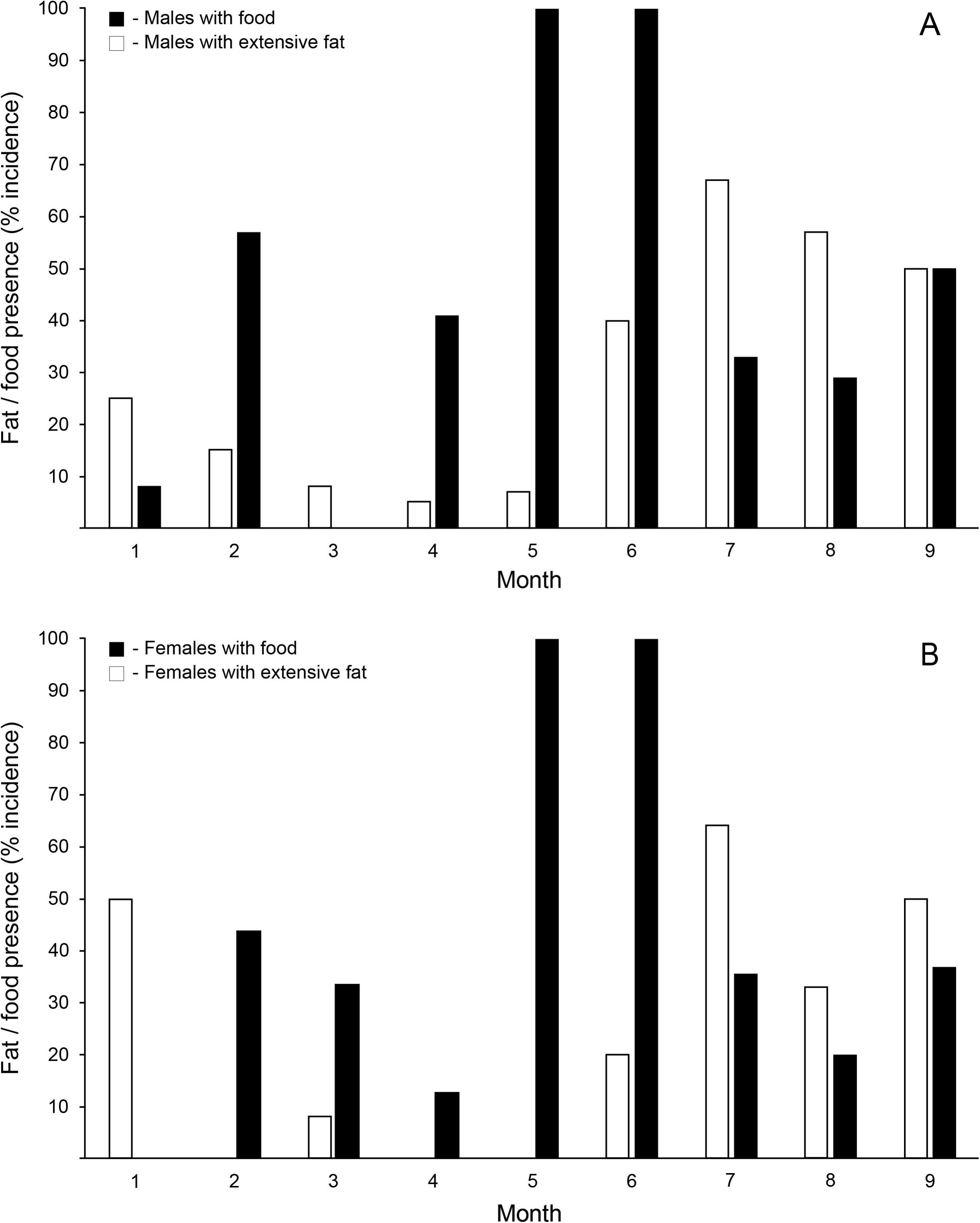
Monthly frequency of extensive fat in the body cavity and the presence of food in the stomach for 104 male (A) and 98 female (B) Rio Grande leopard frogs (*Lithobates berlandieri*) from Texas (USA).

From January–September, the monthly percentage of females with extensive fat development was lowest during February–May, and gradually increased to a peak in July and was relatively high thereafter (Figure 5B). The monthly percentage of females containing food was lowest in January (0%) and increased rapidly to 100% of the females in May and June, and then declined to about 30% during July–September (Figure 5B).

### Ovarian cycle

Gravid females (ovarian stage 4) were detected in each month during January–September (Figure 6). Most gravid females were found in January–June and the fewest in August and September. The months with the lowest frequency of stage 4 females also exhibited the greatest frequency of stage 1 females (July–September), indicating that oviposition is generally finished by late□summer and that oogenesis for oviposition during the upcoming winter had begun. To that end, egg masses have been collected 5 January 1978 from Presidio County (UTEP 4303), and larvae on 3 January 1978 from Presidio County (UTEP 4302) and 21 June 1980 from Kendall County (UTEP 10073).

**Figure 6.**
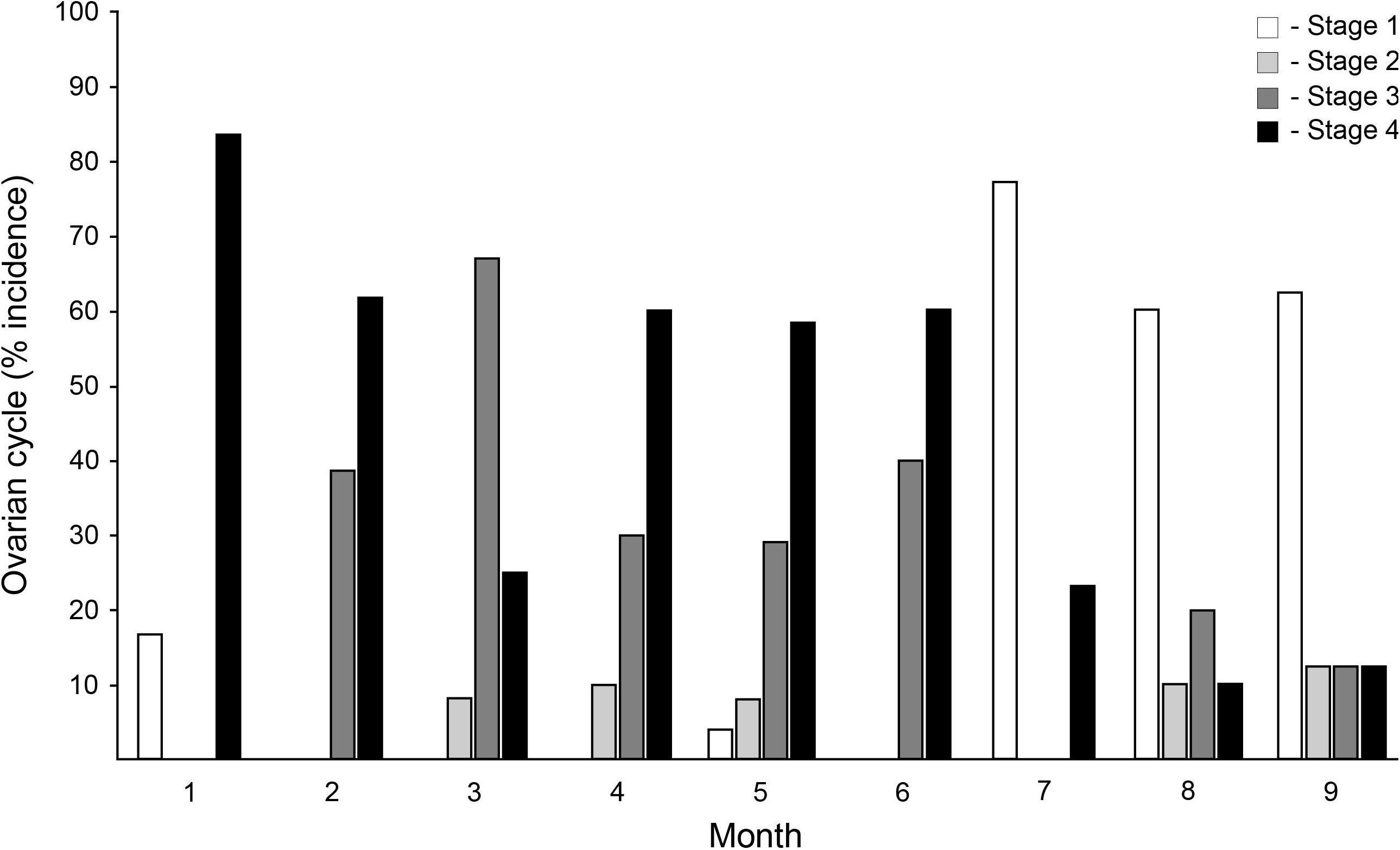
Monthly ovarian cycle of 101 female Rio Grande leopard frogs (*Lithobates berlandieri*) from Texas (USA). See text for details on reproductive stages.

### Clutch and egg size

The mean clutch size of 15 gravid females (SVL = 82.73 ± 14.38 mm; range 59.6–102.8 mm) was estimated at 3,107 ± 2,003.25 eggs (range = 643–8,288 eggs). The body sizes for the 15 females measured for clutch sizes were normally distributed (Shapiro-Wilk test: W = 0.9389, *P* = 0.369), yet their clutch sizes were not (Shapiro-Wilk test: W = 0.86077, *P* = 0.025). Clutch size was positively correlated with body size (Spearman’s test: S = 170, rho = 0.696, *P* = 0.0052; Kendall’s test: T = 80, tau = 0.5238, *P* = 0.0059; Pearson’s test: R^2^ = 0.44, *P* = 0.0068) (Figure 7). The mean ovum diameter from 150 measured ova from 15 gravid females was 1.31 ± 0.27 mm (range = 0.9–2.0 mm). Maximum ovum size was not correlated with either body size (R^2^ = 0.004, F = 0.05, *P* = 0.82) or clutch size (R^2^ = 0.0003, F = 0.003, *P* = 0.96).

**Figure 7.**
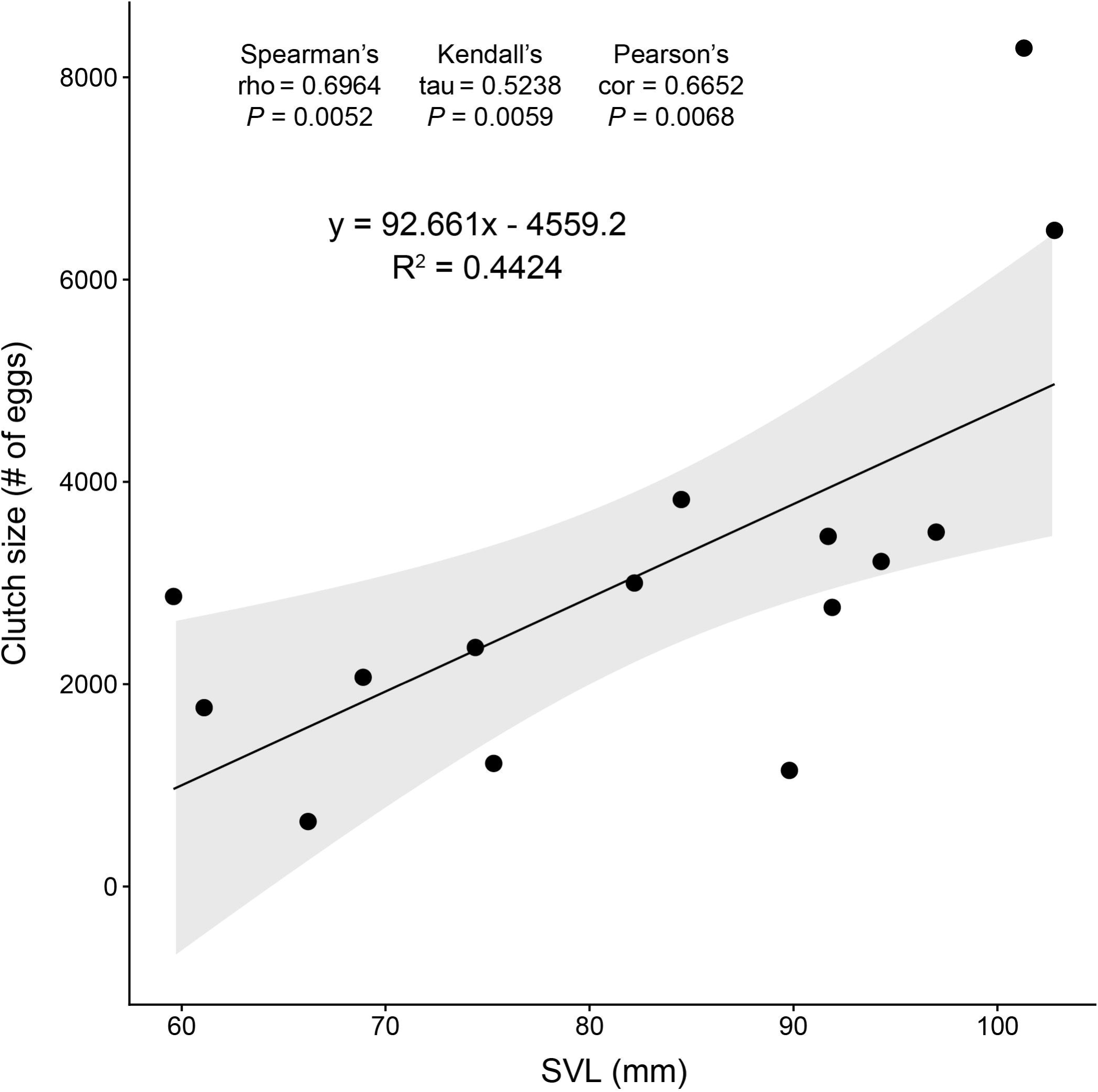
The relationship between clutch size and body size (SVL) in 15 female Rio Grande leopard frogs (*Lithobates berlandieri*) from Texas (USA). Gray shading indicates a 95% confidence interval.

### Fat and food periodicity per ovarian cycle

Nearly 60% of ovarian stage 1 females exhibited extensive fat development (Figure 8). No females in ovarian stage 2 contained extensive fat, less than 20% ovarian stage 3 females contained extensive fat, and less than 10% of females in ovarian stage 4 had this condition. The periodicity of extensive lipid deposits suggests that the depletion of fat stores coincided with egg and clutch development. The monthly distribution of females with extensive fat corroborates a deceased occurrence of fat stores during the probable egg-laying season (Figure 5B). The frequency of ovarian stage 1 females containing food was lowest (21%) (Figure 8). More than half of the females in ovarian stages 2 (60%), 3 (54%), and 4 (58%) all contained food. The peak months of females with food (100% in May and June) suggests that widespread feeding commenced at the end of a spring egg-laying bout (Figure 5B).

**Figure 8.**
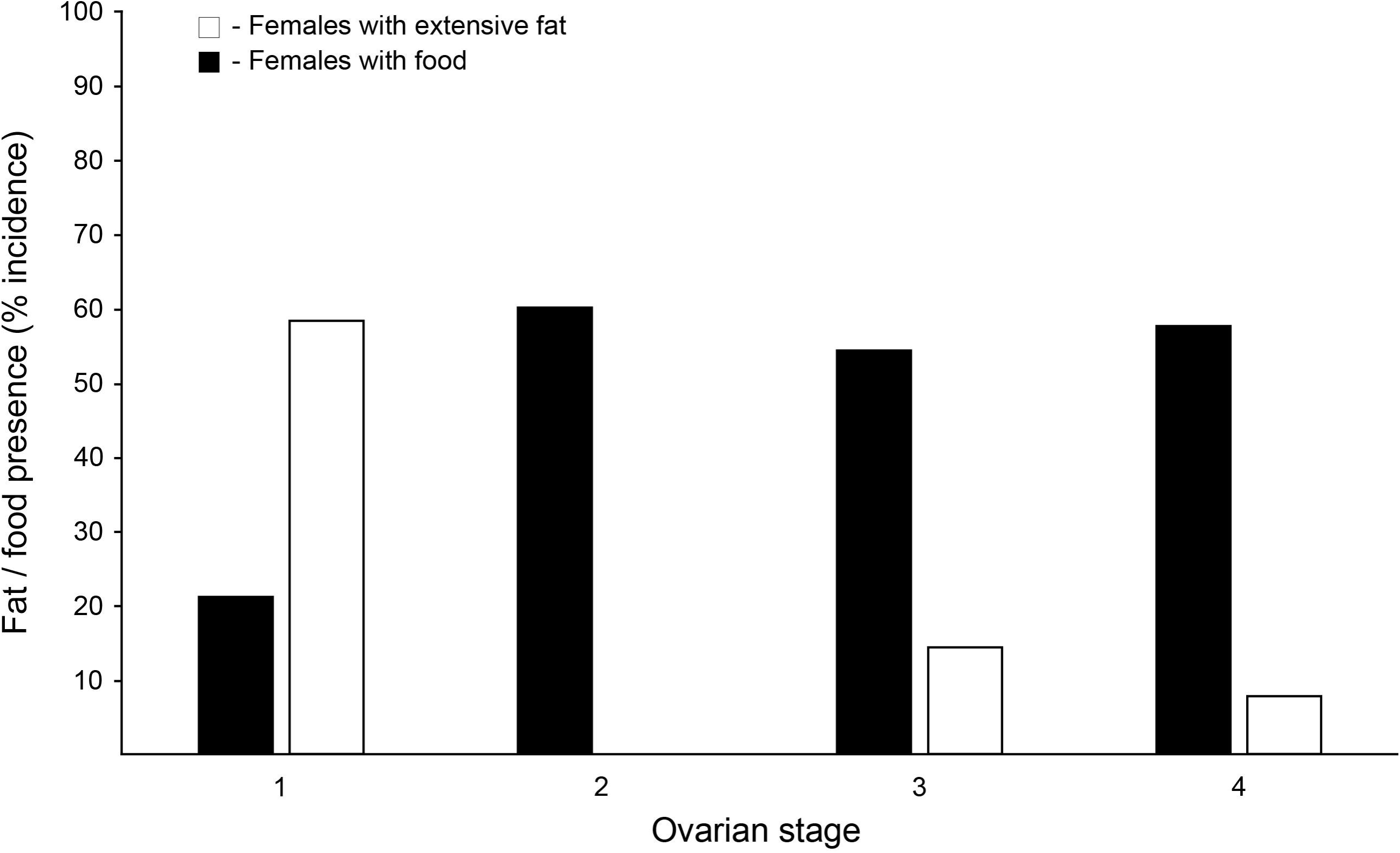
Frequency of extensive fat and the presence of food in each of the four ovarian stages of 98 female Rio Grande leopard frogs (*Lithobates berlandieri*) from Texas (USA). See text for details on reproductive stages.

### Growth and sexual maturity

The smallest frog that had four legs and a tail (i.e., metamorphosling) (SVL = 19.5 mm) was collected on 11 June 1978 in Kimble County (USNM 214057) (Figure 9). Metamorphoslings found after June included an individual (SVL = 23.3 mm) collected on 23 July 1960 in Bexar County (USNM 157194), two individuals (SVL = 24.3 mm and 25.6 mm) collected on 6 August 1960 in Bexar County (USNM 157198–99), and an individual (SVL = 30 mm) collected on 10 September 1941 in Hays County (CM 21140). The smallest juvenile (SVL = 18.6 mm) was collected on 6 August 1960 in Bexar County (USNM 157204). Individuals smaller than 35 mm collected before June were uncommon and included a well-developed tadpole with four legs and a tail (SVL = 28.6 mm) collected on 17 January 1987 in Terrell County (CM 115843), a juvenile (SVL = 33.2 mm) collected on 17 February 1935 in Bexar County (CM 9611), and a frog with its tail (SVL = 30.2 mm) collected on 2 May 1931 in Atascosa County (CM 18424).

**Figure 9.**
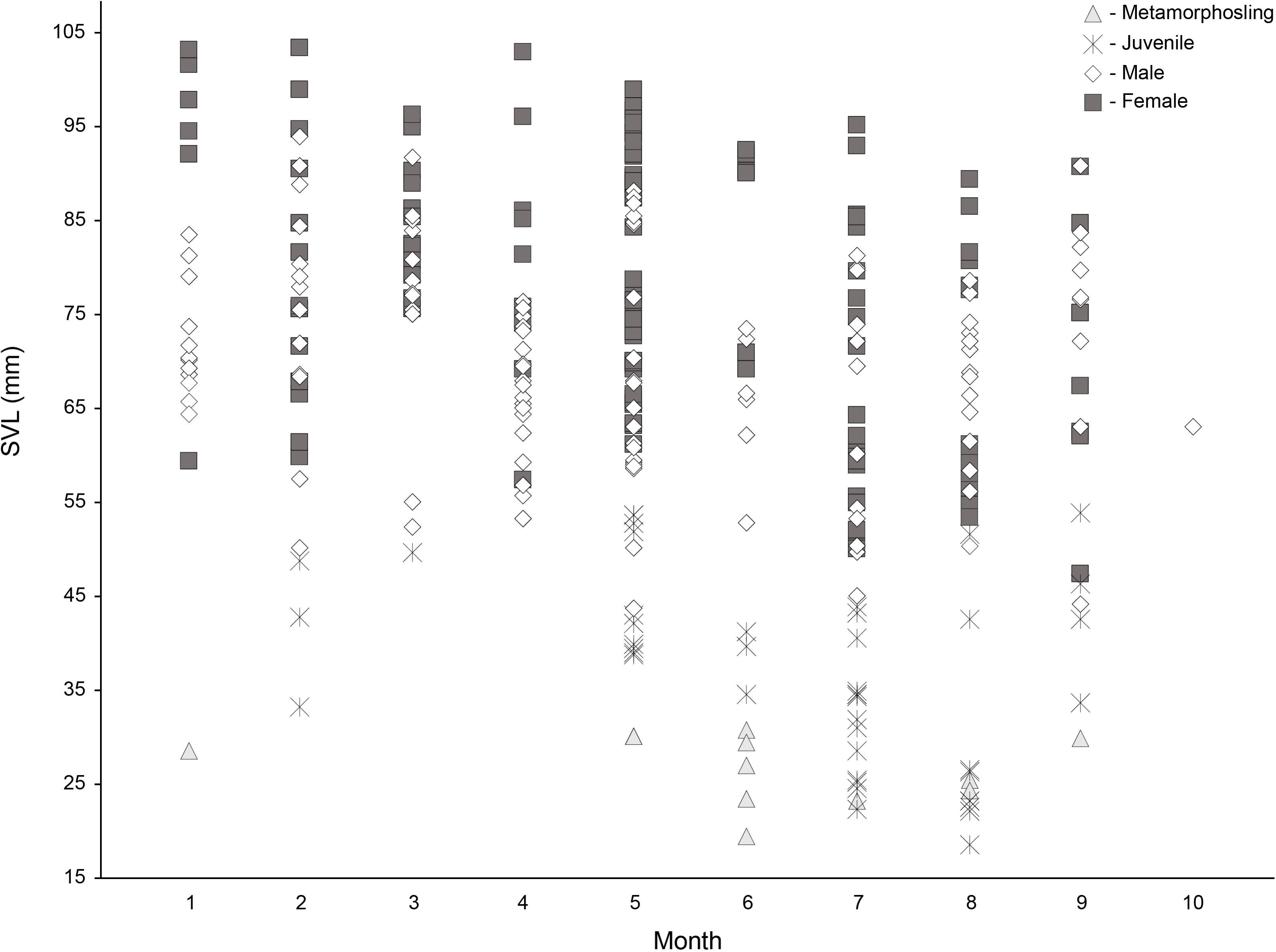
Monthly distribution of body sizes (SVL) of 126 males, 128 females, 44 juveniles, and 11 metamorphoslings (four legs with a tail) of the Rio Grande leopard frog (*Lithobates berlandieri*) from Texas (USA).

The smallest adult male was 43.6 mm in SVL and collected on 21 May 1970 in Irion County (USNM 225484). The smallest apparently sexually active male (based on enlarged testes and thumbs) was collected on 13 February 1970 in Brazos County (UTEP 8606, SVL = 50.1 mm). The smallest gravid female (stage 4) was 57.2 mm in SVL and collected on 3 April 1971 in Coke County (USNM 225450), the smallest ovarian stage 3 female was 70.8 in mm and collected on 11 June 1973 in Jeff Davis County (CM 57860), the smallest ovarian stage 2 female was 60.9 in mm and collected on 13 May 1907 in Cameron County (CM 2257), and the smallest ovarian stage 1 female was 47.3 in mm and collected on 19 September 1968 in Presidio County (USNM 225524). The mean SVL of ovarian stage 1 females was 59.64 ± 7.23 mm (range 47.3–75 mm; n = 23). We note that the smallest ovarian stage 1 females could have been still maturing with egg laying likely occurring at larger sizes/older age classes. The mean SVL of ovarian stage 2 females was 74.73 ± 10.97 mm (range 60.9–86 mm; n = 6), ovarian stage 3 females was 85.94 ± 7.67 mm (range 70.8–98.6 mm; n = 28), and ovarian stage 4 females was 84.59 ± 12.63 mm (range 57.2–103.2 mm; n = 44). Mean body size of gravid females (stage 4) was significantly larger than all other ovarian stages combined (non-gravid female stages 1–3 [74.15 ± 14.69 mm, range 47.3–98.6 mm, n = 57) (t = 3.76, df = 99, *P* = 0.0001).

Tadpole transformation was likely 2–4 months, as evidenced by the seasonal distribution of body sizes (Figure 9). At approximately 3 mm growth per month, males reached a minimum body size at sexual maturity (50. 1 mm) within 3–4 months after metamorphosis and mean adult body size (69.5 mm) almost 14 months post-metamorphic age. Females reached a minimum body size at sexual maturity (57.2 mm) about 6–7 months after metamorphosis and mean adult body size (77.5 mm) almost 17 months post-metamorphic age.

## Discussion

The Rio Grande leopard frog occupies aquatic habitats across a wide range of latitudes in North America. In Texas, we found that *L*. *berlandieri* conforms to general patterns of reproduction in leopard frog species from temperate and sub-tropical latitudes. Nevertheless, a range-wide synthesis of life history in *L*. *berlandieri* has been complicated by equivocal distributional boundaries in México and a general lack of descriptive data for various reproductive traits. We will discuss our findings from Texas in the context of data from elsewhere in the state and from some populations in México—given the known limitations—as an attempt to provide an overview of reproduction in *L*. *berlandieri*.

The reproductive season for the Rio Grande leopard frog in Texas is relatively extended compared to other North American leopard frogs, yet overall patterns were not entirely dissimilar to congeners (e.g., Meshaka et al. 2009; Meshaka and Hughes 2014; Hughes et al. 2017). *Lithobates berlandieri* has been heard calling year-round in Texas (Blair 1961; Hillis 1981; Tipton et al. 2012; Dodd 2013), but calling does not always translate to breeding. From Tornillo Creek in Big Bend National Park, Minton (1958) observed several reproductive activities during the late-winter and spring: Egg masses alongside large and small tadpoles on 16 February; juvenile frogs, new egg masses, and calling males on 25 April; and few frogs of any size after heavy rains on 10 May. From the Rosillos Mountains, also in Big Bend, Minton (1958) detected similar patterns of reproductive activities: New egg masses from 20 February to 15 March; well-developed tadpoles and additional new egg masses on 6 April; and further north in Brewster County, he found egg masses and calling males on 21 July. Blair’s (1961) report from Travis County during 1955–1959 is by far the most detailed treatment of seasonal reproduction in *L*. *berlandieri* prior to our study. Blair (1961) reported winter calling, stimulated by adequate temperature and recent rains, which began as early as 15 December in 1957 to as late as 14 February in 1956, and summer calling that only ceased when the pond dried, yet he found a bimodal pattern to the breeding season. Blair (1961) summarized the timing of bimodal breeding in *L*. *berlandieri* as a winter-spring bout that extends from 8 February to 13 May and an early fall bout from 22 July to 14 October based on recorded egg masses (n = 23), amplexing pairs (n = 19), and gravid females (n = 32) observed during 15–28 February, 12 April–9 May, and 19 July–10 October. Blair’s (1961) fall breeding bout was slightly greater in intensity that the winter-spring bout (55 reproductive incidents to 43). Dayton et al. (2007) reported breeding to be nearly year-round from Brewster County, with observed records of calling males and egg masses in January, March, May–October, and December, yet they did not distinguish if egg masses were found in each of those months. Hillis (1981) found that the breeding season occurred in the fall and the late-winter–spring from Llano and Gillespie Counties, yet only during fall–early winter when sympatric with other leopard frog species. In New Mexico, *L*. *berlandieri* breeding is thought to peak in the spring with calling heard during March–August, and egg masses have been found as early as 22 March yet most were detected from April to early July (Degenhardt et al. 1996). From Eddy County, New Mexico, Scott and Jennings (1985) reported eggs, variously sized tadpoles, and metamorphoslings (Gosner stages 40–42) in April and August, and tadpoles of various sizes in June, October, and November. From introduced populations in Arizona, breeding is assumed to occur all year because calling has been heard from February–October and pairs observed in amplexus as late as October (Rorabaugh 2005). The breeding season for many populations in México has not been characterized, but it is assumed to occur throughout the year (Campbell 1998). From a small pond in the highlands of Zacatecas, México, Drummond and Garcia (1989) studied predator-prey dynamics between garter snakes (*Thamnophis* spp.) and *L*. *berlandieri*, for which they found that tadpoles were present during February–September and tadpole abundance was subject to seasonal variation but exhibited a peak in March and slowly declined thereafter during both 1982 and 1983. Late-stage tadpoles (Gosner stages 39–45), however, displayed a distinct peak in abundance during June of both sampling years and adult frogs were most abundant during May and June in 1982 (Drummond and Garcia 1989). Based on these collective seasonal breeding observations, a general consensus has emerged: Breeding occurs in the fall and continues throughout the winter and spring until mid-summer with salient major breeding bouts in early fall and early spring in most populations across the species range. It seems likely that breeding in *L*. *berlandieri* in Texas is only interrupted by the seasonal extremes in temperature that would occur in the winter and summer, yet in years and/or areas where temperatures are maintained within an appropriate range (11–32°C [Blair 1961]) breeding could occur all year if stimulated by adequate precipitation.

Development of the relatively large *L*. *berlandieri* tadpoles can be prolonged in Texas depending on whether eggs are laid in the winter/spring or in the fall. In McLennan County, Hillis (1982) found that the *L*. *berlandieri* tadpoles (SVL = 27.2 ± 3.1 mm, range 19.7–31.5 mm, n = 25) were larger than the tadpoles of sympatric leopard frog species and that they overwintered. In Brewster County, Dayton et al. (2007) suggested that the larval period is 4–9 months and that the local conditions of specific breeding ponds, especially water temperature and hydroperiod, strongly influence larval-period dynamics. In Arizona, tadpoles have been observed to overwinter as well (Rorabaugh 2005). Drummond and Garcia (1989) suggested that tadpoles developed from 10 to 17 weeks in Zacatecas (México), which is a more rapid larval development time than observed in Texas (Dayton et al. 2007; this study). From northern Guatemala, the Yucatán Peninsula, and Belize, Campbell (1998) found tadpoles during February–September, and metamorphs (or juveniles) in March and April.

Adult body sizes of *L*. *berlandieri* in Texas were large and generally similar to reported sizes for congeners in México and in the United States (Campbell 1998; Lemos-Espinal and Smith 2007; Dodd 2013). The degree of SSD in *L*. *berlandieri* was consistent with other North American leopard frogs (Dodd, 2013), including *L*. *sphenocephalus* (Hughes et al. 2017). In New Mexico, *L*. *berlandieri* males (mean SVL = 64.4 mm) were almost 10 mm smaller than females (mean SVL = 73.5 mm) (Degenhardt et al. 1996). Our sample means were nearly 5 mm larger than both the males and females from New Mexico and also displayed a lesser degree of SSD. In the United States, the maximum body size for the species was a recorded from a female (SVL = 114 mm) of an introduced population in Arizona (Brennan and Holycross 2006), which is >10 mm larger than the largest specimen we examined—a female collected on 26 February 1971 in Uvalde County with SVL = 103.2 mm (USNM 225660). Dayton et al. (2007) indicated that the average body size of *L*. *berlandieri* in Big Bend National Park ranges 63.5–101.6 mm. Powell et al. (2016) reports a range of 57–100 mm for body sizes of *L*. *berlandieri* in the United States. Data on body sizes from elsewhere in the species distribution are uncommon and from its southern range limit can be misleading. From northern Guatemala, the Yucatán Peninsula, and Belize, Campbell (1998) reported that *L*. *berlandieri* males reach body sizes of 75–85 mm in SVL and females are much larger with a range of 115–120 mm in SVL. However, Campbell (1998) suggested that the species identity of leopard frogs in southern México and Guatemala is unresolved and that frogs assigned to this species may represent several species. Zaldívar-Riverón et al. (2004) found that leopard frog populations along the Atlantic coast immediately south of Veracruz to the Yucatán Peninsula and northern Guatemala are in fact *L*. *brownorum* and on the Pacific coast, *L*. *forreri*. Consequently, descriptive data reported from populations south of Veracruz may in fact represent a different species (e.g., Luque-Montes et al. 2018).

The increase in feeding shortly after the spring breeding bout is consistent with Parker and Goldstein’s’ (2004) findings of an increase in stomach contents in a fall versus spring sample of *L*. *berlandieri* from Texas. An increase in fat deposits after June and a corresponding increase in the frequency of stage 1 females—which are either spent or still maturing if very small—beginning in July were indicative of a mid-summer breeding hiatus as the aquatic breeding habitats likely dried because of high temperatures. Nevertheless, records of females beginning oogenesis (stage 2 and 3) increased shortly thereafter for a likely breeding bout in the early fall. Patterns of ovarian cycling related to oogenesis, lipid deposition, and food intake in *L*. *berlandieri* from Texas are generally consistent with the available data for other North American leopard frog species (e.g., Dodd 2013; Meshaka and Layne 2015). However, yet to be examined are populations from elsewhere in the species range, the closely related coastal leopard frog species in México (Zaldívar-Riverón et al. 2004), and many of the 27 unique lineages of the subgenus *Pantherana* (which includes *L*. *berlandieri*) identified by Yuan et al. (2016).

We provided the first estimates of clutch size and egg size for *L*. *berlandieri* and thus our data fill-in other conspicuous knowledge gaps for an apparently well-studied North American frog (*sensu* Mitchell and Pague 2014). Further, we described the first evidence of geographic variation in adult body sizes from New Mexico to Texas, which suggested that populations at higher latitudes possess smaller body sizes. Detailed examinations into other life-history characters from populations in New Mexico and greater sampling across latitude in general would provide the material necessary to quantify the extent that geography influences other aspects of reproduction in *L*. *berlandieri*. Of the leopard frog species examined, many exhibit latitudinal signatures in life-history characteristics (Morrison and Hero 2003). Descriptive studies into life-history variation within and across species may lead to the discovery of general patterns of anuran reproduction. Indeed, the significance of descriptive data in the study of life-history evolution in amphibians has long been recognized (Noble 1927), yet the collection of reproductive data has lagged far behind that of other sources, such as molecular. We can infer general patterns of trait evolution by mapping data onto phylogenies (Blackburn 2008), and while the appropriate framework exists (e.g., Yuan et al. 2016), the dearth of descriptive data available for leopard frogs impedes a robust examination into the evolution of their life history. To that end, our study on the life history of the Rio Grande leopard frog has not only provided the descriptive data necessary to more rigorously explore the impacts that future environmental changes may have on its reproduction, but our data also represent a contribution towards better understanding the life-history evolution in Ranidae.

## Acknowledgements

We are tremendously grateful to Carl S. Lieb, Teresa Mayfield, Eli Greenbaum (UTEP Biodiversity Collections), Stephen P. Rogers (Carnegie Museum of Natural History), Kenneth Tighe, Addison Wynn, and Kevin de Queiroz (United States National Museum) for unfettered access to specimens, quiet places to work, and excellent company during this research project.

## ORCID

Daniel F. Hughes https://orcid.org/0000-0001-5883-5324

